# Inducible CRISPR activation screen for interferon-stimulated genes identifies OAS1 as a SARS-CoV-2 restriction factor

**DOI:** 10.1101/2021.09.22.461286

**Authors:** Oded Danziger, Roosheel S Patel, Emma J DeGrace, Mikaela R Rosen, Brad R Rosenberg

## Abstract

Interferons establish an antiviral state in responding cells through the induction of hundreds of interferon-stimulated genes (ISGs). ISGs antagonize viral pathogens directly through diverse mechanisms acting at different stages of viral life cycles, and indirectly by modulating cell cycle and promoting programmed cell death. The mechanisms of action and viral specificities for most ISGs remain incompletely understood. To enable the high throughput interrogation of ISG antiviral functions in pooled genetic screens while mitigating the potentially confounding effects of endogenous IFN and potential antiproliferative/proapoptotic ISG activities, we adapted a CRISPR-activation (CRISPRa) system for inducible ISG induction in isogenic cell lines with and without the capacity to respond to IFN. Engineered CRISPRa cell lines demonstrated inducible, robust, and specific gRNA-directed expression of ISGs, which are functional in restricting viral infection. Using this platform, we screened for ISGs that restrict SARS-CoV-2, the causative agent of the COVID-19 pandemic. Results included ISGs previously described to restrict SARS-CoV-2 as well as multiple novel candidate antiviral factors. We validated a subset of candidate hits by complementary targeted CRISPRa and ectopic cDNA expression infection experiments, which, among other hits, confirmed OAS1 as a SARS-CoV-2 restriction factor. OAS1 exhibited strong antiviral effects against SARS-CoV-2, and these effects required OAS1 catalytic activity. These studies demonstrate a robust, high-throughput approach to assess antiviral functions within the ISG repertoire, exemplified by the identification of multiple novel SARS-CoV-2 restriction factors.

## Introduction

Interferons (IFN) act as key mediators of the host response to viral pathogens by establishing a general antiviral state through the coordinated induction of hundreds of interferon stimulated genes (ISGs) in infected and uninfected “bystander” cells^1^. ISGs encode functionally diverse gene products, including antiviral effectors that antagonize distinct steps of viral life cycles^2^, but the antiviral mechanisms of most individual ISGs remain unknown. Studies that have systematically characterized the effects of single ISGs have demonstrated that a limited number of individual ISGs, primarily transcription factors and DNA/RNA sensors, can broadly restrict infection by multiple viruses upon overexpression in target cells^3–7^. Other individual ISGs have been found to restrict or even to enhance the replication of specific viruses. In addition to direct antiviral effectors, the ISG repertoire also includes genes that induce antiproliferative and/or proapoptotic programs in response to viral infection or DNA damage, thereby limiting viral spread and impeding oncogenesis^8–11^.

A robust IFN response is critical for host defense against novel respiratory viruses to which immune memory from prior exposure has not been established^12^. As such, multiple lines of evidence have implicated IFN as a key component of the host response to Severe Acute Respiratory Syndrome coronavirus 2 (SARS-CoV-2), the etiological agent of COVID-19. Although IFN can effectively block SARS-CoV-2 infection *in vitro*^13–16^, the activity of IFN in different physiological contexts is more complex. Characterizations of the IFN response to SARS-CoV-2 infection in a variety of model systems suggest that SARS-CoV-2 infection may not elicit robust IFN production and ISG expression, but can induce high levels of proinflammatory cytokines^17^. These findings are consistent with single cell RNA sequencing (scRNA-seq) immune profiling studies that have reported inflammatory gene signatures and less robust IFN/ISG expression in immune cells from individuals infected with COVID-19 as compared to individuals infected with Influenza^18^. However, transcriptomic analyses of bronchoalveolar lavage fluid (BALF) from COVID-19 patients detected a strong ISG signature along with proinflammatory cytokine gene expression in immune cells^19^. Dysregulation of the IFN response in SARS-CoV-2 infection can be driven by both direct viral antagonism of innate immune mechanisms, as well as by host characteristics such as age, genetics and other comorbidities^20^. Additionally, several studies have identified autoantibodies against IFN as a significant negative survival factor for severe COVID-19, further emphasizing the prominent role of IFN in SARS-CoV-2 pathogenesis and clearance^21–24^. Importantly, IFN-mediated effects may also be detrimental to COVID-19 outcomes. For example, disruption of the lung epithelium by type III IFN-dependent processes has been hypothesized to expose patients to secondary infections by opportunistic bacteria^25^. Taken together, while IFN and the downstream expression of ISGs can functionally restrict SARS-CoV-2, the site(s), cell types, amount, and timing of IFN production and response play critical roles in COVID-19 pathogenesis and outcomes.

The specific mechanisms by which IFN restricts SARS-CoV-2 have not been fully characterized. To date, of the hundreds of ISGs, only a handful have been found to restrict SARS-CoV-2 infection in different *in vitro* systems. Lymphocyte antigen 6 complex, locus-E (LY6E) was identified as a SARS-CoV-2 restriction factor in ISG ectopic cDNA expression screens^26^. A transposon-mediated screen identified the MHC-II invariant chain CD74 as a block to SARS-CoV-2 entry^27^. In addition, overexpression of BST2 (encoding Tetherin), an anti-HIV effector that prevents the release of nascent virions^28^, has also been found to restrict SARS-CoV-2 by impeding virion release^29^. TRIM25, an interferon-induced E3 ubiquitin ligase that enhances antiviral responses downstream of RIG-I, has been shown to interact specifically with SARS-CoV-2 RNA and thereby reduce infection^30^.

Studies evaluating the antiviral potential of individual ISGs are often implemented through the ectopic expression of ISG cDNA libraries followed by viral challenge to screen for those ISGs that confer resistance^3,4,29,31,32^. While effective in identifying the antiviral potential of many ISGs, arrayed ISG cDNA screens are not without drawbacks including limited throughput, technically demanding cloning and validation of individual expression constructs, and high costs. Ectopic cDNA overexpression can be prone to artifactual expression patterns or functions^33,34^. In addition, cDNA expression screen libraries typically include only one isoform per gene, and therefore may overlook isoform-specific antiviral activities, as have been described for many ISGs^27,35–39^.

CRISPR-activation (CRISPRa), in which guide RNA (gRNA)-directed endonuclease-deficient Cas9 along with transcriptional activators are targeted to a gene of interest (GOI) to induce expression, offer an alternative to cDNA ectopic expression screens. Advantages include easy to produce gRNA libraries, physiologically relevant expression levels, and multiplexing capabilities^40^. Gene transcription is initiated from endogenous promoters, enabling the expression of multiple gene isoforms. Studies in which genome-wide libraries of activating gRNAs uncovered important host factors in viral infection systems provide an important proof of concept to the characterization of ISGs using CRISPRa^41,42^. Importantly, two recent preprints report genome-wide CRISPRa screens for genes with antiviral potential against SARS-CoV-2^43,44^.

Here, we report an ISG-focused CRISPRa screen to identify ISGs that modulate SARS-CoV-2 infection in lung epithelial cells. To mitigate the potential antiproliferative and/or proapoptotic effects of certain ISGs that could impact their library representation, we engineered a Doxycycline (Dox)-inducible CRISPRa system that enables precise temporal control of ISG induction. Using a pooled screen strategy, we tested more than 400 ISGs for effects on SARS-CoV-2 infection in both wildtype cells and isogenic cells engineered to be insensitive to IFN. High ranking antiviral ISG hits included SARS-CoV-2 restriction factors previously identified in recent screens (*LY6E*^26^, *CD74*^27^, *TRIM25*^30^ *and ERLIN1*^29^). We identified and validated antiviral roles for additional ISGs such as *CTSS* (Cathepsin S). We also identified *OAS1* (2′-5′-oligoadenylate synthetase 1) as a SARS-CoV-2 restriction factor capable of inhibiting viral infection and the generation of progeny virus. Taken together, our findings demonstrate the utility of a novel inducible CRISPRa platform for antiviral genetic screens and identify multiple ISGs capable of restricting SARS-CoV-2.

## Results

### Inducible CRISPRa system in A549 cells

We developed an optimized platform for pooled, positive selection ISG screens by adapting the well-established SunTag CRISPRa technology^45^, and engineering it into A549 lung adenocarcinoma cell lines, a widely employed model for respiratory virus infection^46–48^. First, we reasoned that the antiproliferative and/or proapoptotic properties of some ISGs^49^ could affect their relative representation in pooled libraries prior to infection experiments and/or independent of potential effects on virus susceptibility. To mitigate these effects, we reengineered the transcriptional transactivator component construct of the SunTag CRISPRa system to allow for Doxycycline (Dox)-inducible expression, and thereby Dox-inducible regulation of gRNA-targeted gene expression (Fig. 1A). Next, we expected that IFN secretion in response to infection, with corresponding autocrine and paracrine induction of broad ISG expression throughout cultures, could interfere with assessments of individual CRISPRa-induced ISG effects within different cells in pooled screen experiments. Therefore, we transduced the modified SunTag components into A549^ΔSTAT1^ cell lines defective in their capacity to respond to IFN (A549^ΔSTAT1^-SunTag^50^), as well the wildtype A549 cell line (A549-SunTag, intact IFN response) from which they were derived. After selecting and expanding dual antibiotic-resistant single cell clones, we evaluated their capacity for Dox-inducible, guide RNA (gRNA)-targeted gene expression. We transduced A549-SunTag and A549^ΔSTAT1^-SunTag cell lines with lentiviral constructs expressing gRNAs targeting the promoter region of *MX1*, a well-characterized ISG that restricts multiple viruses^51,52^, as well as with non-targeting gRNA (NTG) controls. Following puromycin selection of gRNA-transduced cells, cultures were treated with Dox for 48 hours, and *MX1* mRNA expression was assessed by quantitative RT-PCR (qRT-PCR). In both A549-SunTag and A549^ΔSTAT1^-SunTag genotypes, Dox induced robust increases in *MX1* mRNA levels in cells expressing *MX1* gRNAs (Fig. 1B), while Dox treatment of cells expressing NTG or non-transduced cells exhibited minimal changes in *MX1* gene expression. Comparison to cells pretreated with IFNα2b indicated that CRISPRa induced *MX1* mRNA expression to similar levels (Fig. 1B). Interestingly, baseline levels of *MX1* mRNA exhibited higher Ct values in A549^ΔSTAT1^ cells compared to their wildtype counterparts, resulting in greater fold change values for *MX1* expression upon Dox treatment (Fig. 1B-C). This suggests that, even in the absence of exogenous IFN stimulation, some ISGs exhibit some level of constitutive STAT1-dependent transcription.

**Figure 1:**
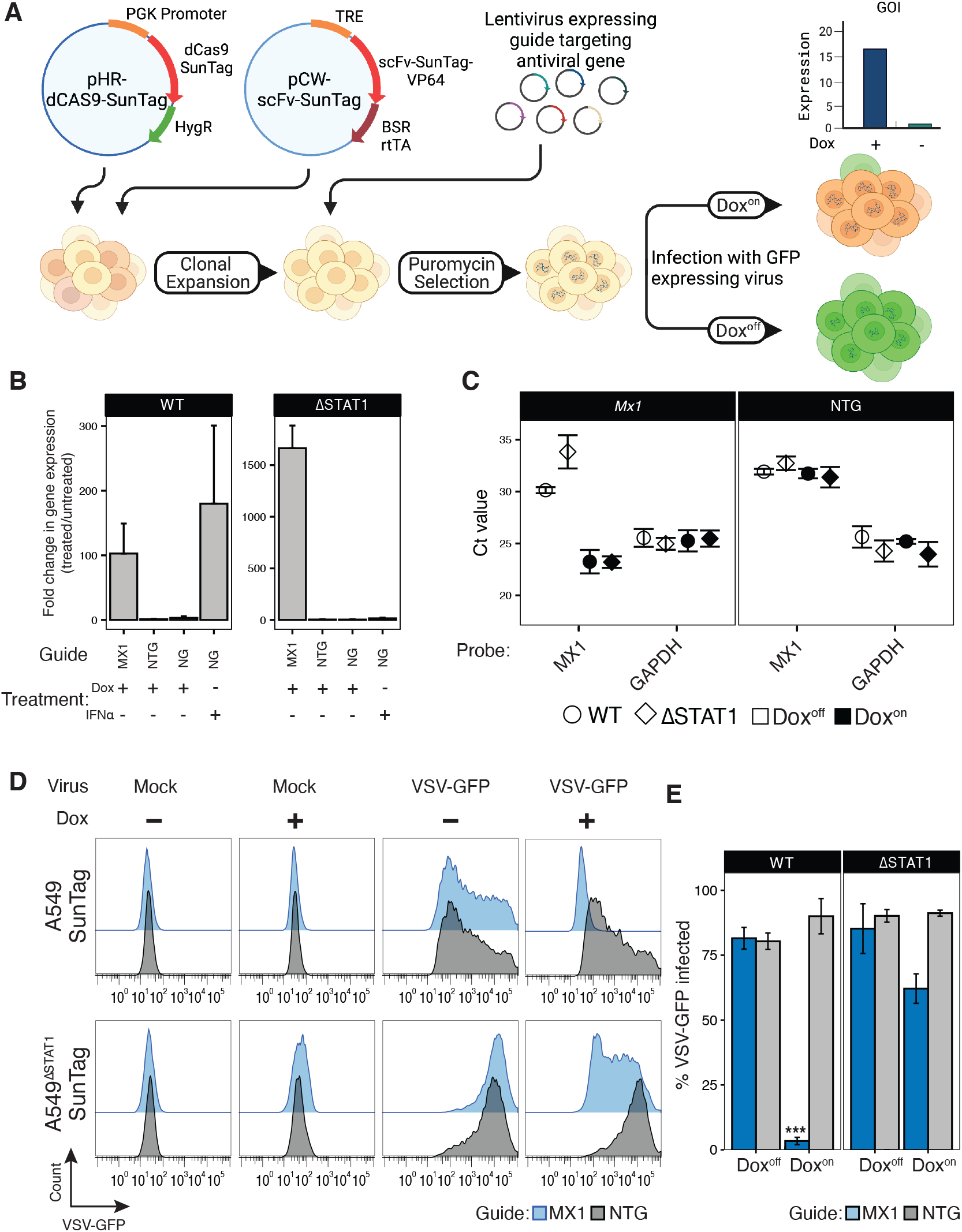
Inducible CRISPRa system demonstrates Dox-regulated, gRNA-specific gene expression and functional block of VSV-GFP infection in isogenic A549 and A549^ΔSTAT1^ cell lines. **(A**) Schematic of the Dox-inducible CRISPRa system for assessing antiviral ISG activities. **(B)** qRT-PCR analysis for *MX1* mRNA in A549-SunTag and A549^ΔSTAT1^-SunTag cells transduced with *MX1* gRNA, non-targeting gRNA (NTG), or in cells with no gRNA (NG), treated with Dox or IFN. Fold change gene expression in Dox^on^ cells relative to Dox^off^ cells calculated using the ΔΔCt method with normalization to *GAPDH*. **(C)** Mean qRT-PCR threshold cycle (Ct) values for *MX1* and *GAPDH* mRNA in A549-SunTag and A549^ΔSTAT1^-SunTag cells transduced with *MX1* gRNA or NTG gRNA (NTG). Error bars indicate ± SD. **(D)** Representative flow cytometry histograms for GFP fluorescence in A549-SunTag and A549^ΔSTAT1^-SunTag transduced with *MX1* gRNA (blue) or NTG (gray) gRNA and infected with VSV-GFP (M.O.I. = 1, 24hr). **(E)** Percent VSV-GFP-positive cells by flow cytometry quantified for n = 3 biological replicates. Bars represent mean ± SD of GFP positive cells across all replicates. Paired ratio Student’s t-test, *** p < 0.0005.

Next, we evaluated the functional antiviral capacity of an ISG in our CRISPRa cells. A549-SunTag and A549^ΔSTAT1^-SunTag cells, transduced and selected to express *MX1* gRNA, were treated with Dox and infected with a GFP-encoding Indiana vesiculovirus (VSV-GFP^53^). Flow cytometry analysis, using GFP expression as a marker for productive infection, demonstrated a near-complete block of infection in Dox^On^ A549-SunTag cells with active *MX1* expression; Dox^Off^ cells and cells expressing an NTG were not protected (Fig. 1D-E). While we also observed a protective effect in A549^ΔSTAT1^-SunTag, a considerable fraction of cells remained susceptible to infection. These results indicate that Dox-inducible, CRISPRa-mediated ISG expression can effectively restrict viral infection. Furthermore, the restriction of VSV-GFP infection by MX1 is enhanced by additional, STAT1-dependent factors likely elicited by IFN production, further highlighting the utility of a STAT1-deficient screening platform for assessing the antiviral activity of individual ISGs. In sum, we have established a functional, Dox-regulated CRISPRa system in isogenic cell lines with either intact or deficient IFN responses that effectively restricts viral infection upon induced expression of antiviral ISGs.

### Inducible CRISPRa ISG screen for SARS-CoV-2 restriction factors

To identify ISGs that restrict SARS-CoV-2, we conducted pooled gene activation screens in our engineered CRISPRa A549-SunTag lines. Our general screening strategy was to evaluate the potential of hundreds of individual ISGs to confer resistance to the cytopathic effects of SARS-CoV-2. We began by introducing ACE2 expression into A549-SunTag and A549^ΔSTAT1^-SunTag cells to enable productive SARS-CoV-2 infection of our CRISPRa cells. Next, we conducted pilot experiments evaluating SARS-CoV-2 cytopathic effect (CPE) for optimization of screen conditions to balance the strength of selective pressure with its duration for robust detection of hits^54^. A549-SunTag ACE2 and A549^ΔSTAT1^-SunTag ACE2 cells were transduced with expression constructs for gRNAs targeting *LY6E*, a known SARS-CoV-2 restriction factor^26^, or NTGs. Following Dox treatment, cultures were infected with SARS-CoV-2 at a range of multiplicities of infection (M.O.I.), plates were fixed every 24 hours, and cell viability was estimated by Methylene blue assay (Supplementary Fig. 1A). Dox^On^ cultures expressing *LY6E* gRNAs exhibited increased viability when infected with SARS-CoV-2. Moreover, A549-SunTag ACE2 cultures exhibited less CPE than A549^ΔSTAT1^-SunTag ACE2 cultures, implicating additional STAT1-dependent antiviral factors. Based on these data, we approximated optimal infection conditions for ISG screens (M.O.I. = 3, harvest at 72 hours post-infection). To construct an ISG gRNA library for screening, we merged lists of ISGs tested in previous studies^3,4^ with a list of genes upregulated by IFNβ treatment in A549 cells^50^ (log_2_ fold-change >2, adjusted p value < 0.05). To focus on antiviral effectors, we excluded known transcription factors^55^, several central Pattern Recognition Receptors (PRRs), and HLA genes (Supplemental table 1 contains the full list of ISGs in the library). For each ISG, we selected 3 gRNA sequences from the optimized Calabrese CRISPRa collection^56^. ISG gRNA sequences were supplemented with an additional 24 NTG controls, for a final list of 1,266 gRNAs targeting 414 ISGs (Supplemental table 2).

Gene activation screens were conducted in both A549-SunTag ACE2 and A549^ΔSTAT1^-SunTag ACE2 cells, in multiple independently transduced clones, across two independent experiments. A549-SunTag ACE2 and A549^ΔSTAT1^-SunTag ACE2 cells were transduced with our ISG gRNA library (M.O.I. = 0.1), puromycin selected, and expanded. 48 hours prior to SARS-CoV-2 infection, appropriate cultures were treated with Dox to induce ISG expression (Fig. 2A). All experiments included Dox^Off^ and Dox^On^ conditions, each of which included Mock or SARS-CoV-2 infection. At 72 hours post-infection, gRNA libraries were prepared from surviving cells and sequenced to assess gRNA relative enrichment/depletion. As we observed somewhat more CPE than expected in the first experiment, we slightly relaxed the selection pressure in the second experiment (additional wash for excess virus, details in *Materials and Methods*). All datasets passed quality control metrics for multiple parameters, including percent read mapping (Supplementary Fig. 1B), and normalized gRNA abundance (Supplementary Fig. 1C). To take advantage of our Dox-inducible system and replicated design for rigorous detection of ISG effects on SARS-CoV-2, we used MAGeCK-MLE^57,58^ to test differential gRNA enrichment with a linear model including factors for Dox treatment (Off/On), SARS-CoV-2 infection status (Mock/Infected), Clone (independent gRNA library transduction 1/2), and Experiment (1/2, Full screen results: Supplemental table 3). Importantly, this analysis strategy enabled assessment of ISG effects on SARS-CoV-2 in the context of potential antiproliferative/proapoptotic ISG effects that might independently alter gRNA abundance (assessed by Dox^On^ Mock *vs* Dox^Off^ Mock and corresponding interaction term in the model).

**Figure 2:**
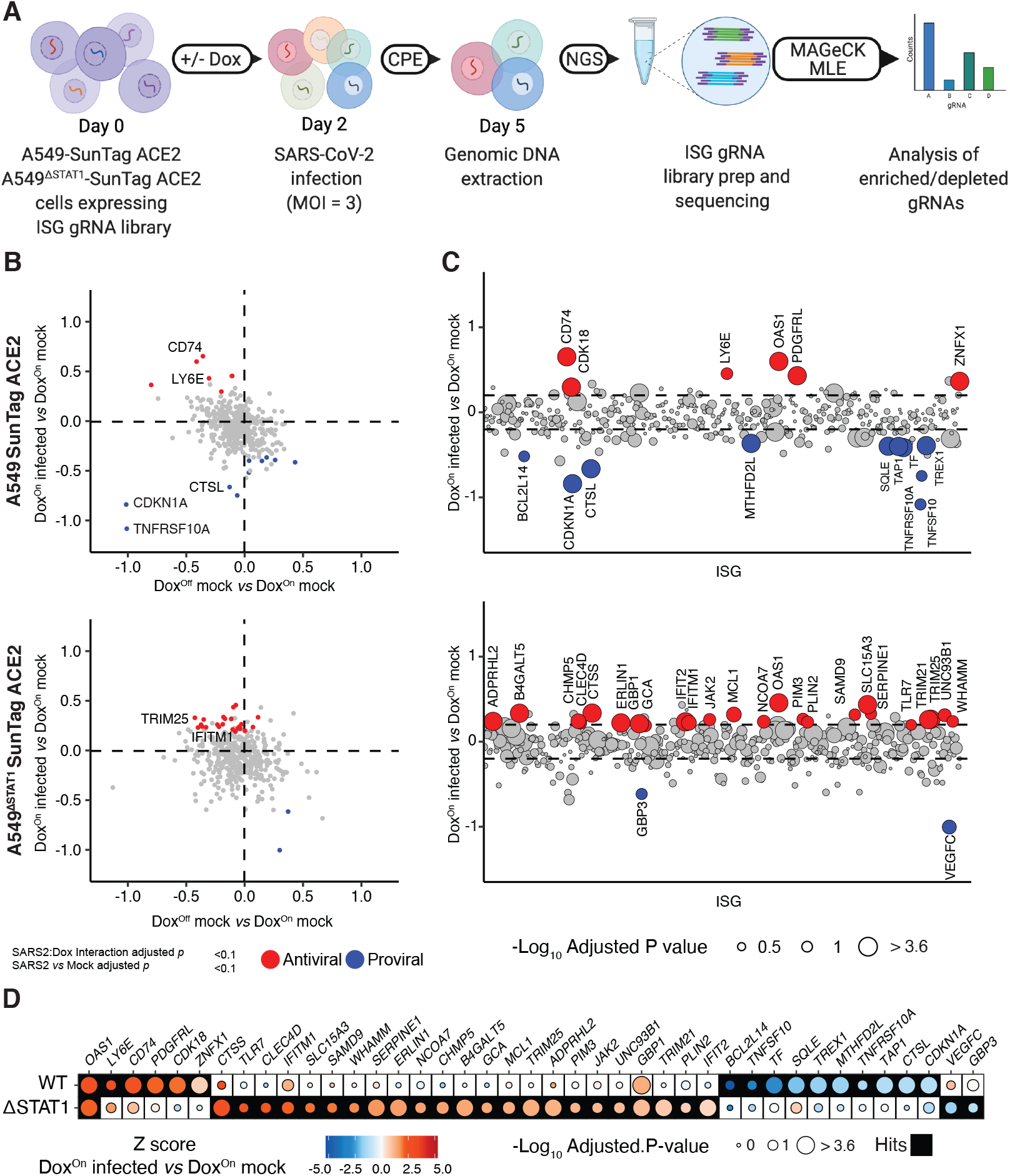
Inducible CRISPRa ISG screen for SARS-CoV-2 restriction factors. **(A)** Schematic of inducible CRISPRa ISG screens for SARS-CoV-2: Single cell clones of A549-SunTag ACE2 and A549^ΔSTAT1^-SunTag ACE2 cells were transduced with a library of 1,266 gRNAs and infected with SARS-CoV-2 (M.O.I. = 3, 72 hours). After onset of SARS-CoV-2 CPE, genomic DNA was extracted from surviving cells, and enriched/depleted gRNAs were evaluated by high throughput sequencing followed by MAGecK MLE analysis. **(B)** MAGeCK-MLE analysis of SARS-CoV-2 ISG screens in A549-SunTag ACE2 and A549^ΔSTAT1^-SunTag ACE2 cells. Plot depicts MAGeCK MLE β scores for Dox status coefficient (Dox^Off^ mock infected *vs* Dox^On^ mock infected, x-axis.), and SARS-CoV-2 infection status coefficient (Dox^On^ SARS-CoV-2 infected *vs* Dox^On^ mock infected, y-axis). ISGs passing significance selection filters (SARS-CoV-2 infection status adjusted p value and SARS-CoV-2:Dox interaction adjusted p value < 0.1) are highlighted in red/blue for “antiviral”/“proviral” effects, respectively. Select subset of annotated SARS-CoV-2 restriction factors and antiproliferative genes are labeled with gene symbols. **(C)** MAGeCK-MLE analysis of SARS-CoV-2 ISG screens in A549-SunTag ACE2 and A549^ΔSTAT1^-SunTag ACE2 cells, aligned by ISG. MAGeCK MLE β scores for SARS-CoV-2 infection status coefficient (Dox^On^ SARS-CoV-2 infected *vs* Dox^On^ mock infected) are plotted for each ISG (x-axis, alphabetical order); dots are sized according to significance of SARS-CoV-2:Dox interaction coefficient (-Log_10_ adjusted p value) and highlighted in red/blue as in (B). Dashed line indicates ± 1 standard deviation of SARS-CoV-2 infection status coefficient β scores. **(D)** Comparison of candidate “antiviral”/“proviral” ISG hits passing significance filters in at least one (A549-SunTag ACE2 or A549^ΔSTAT1^-SunTag ACE2) genotype. Dots are shaded by ISG Z-scores for SARS-CoV-2 infection status coefficient (Dox^On^ SARS-CoV-2 infected *vs* Dox^On^ mock infected), and sized according to the significance of SARS-CoV-2:Dox interaction coefficient (-Log_10_ adjusted p value). Filled boxes indicate ISG passing significance filters for the indicated screen.

To identify ISGs with an effect on SARS-CoV-2 CPE in A549-SunTag ACE2 or A549^ΔSTAT1^-SunTag ACE2 cells, we applied a stringent series of filters for gRNAs differentially enriched by SARS-CoV-2 infection (infection status adjusted p value < 0.1), while accounting for Dox effects (interaction adjusted p value < 0.1). We detected 6 ISGs (A549-SunTag ACE2) and 24 ISGs (A549^ΔSTAT1^-SunTag ACE2) as “antiviral” (i.e. enriched by SARS-CoV-2 infection, Fig. 2B-C). Conversely, we detected 10 ISGs (A549-SunTag ACE2) and two ISGs (A549^ΔSTAT1^-SunTag ACE2) as “proviral” (i.e. depleted by SARS-CoV-2 infection, Fig. 2B-C). Antiviral hits in at least one screen (i.e. wildtype or Δ*STAT1*) included *LY6E, CD74, TRIM25* and *IFITM1* (Fig. 2B-C), each of which has been previously reported to restrict SARS-CoV-2^26,27,30,59^. Proviral hits in at least one *STAT1* genotype included *CTSL* (encoding Cathepsin L), an entry factor for coronaviruses^60^. Additional proviral hits *CDKN1A* (p21) and *TNFRSF10A*, ISGs with antiproliferative and/or proapoptotic effects^61,62^, were depleted upon Dox treatment independently of viral infection, and were further depleted by SARS-CoV-2 infection. This is in line with previous observations that coronaviruses require cell cycle inhibition for optimal replication^63^. Interestingly, when comparing ISG hits between A549-SunTag ACE2 and A549^ΔSTAT1^-SunTag ACE2 screens (Fig. 2D), we found only a single common antiviral hit across genotypes: *OAS1* (encoding 2′-5′-oligoadenylate synthetase 1). Not surprisingly, our results suggest that STAT1-dependent transcription, perhaps in response to endogenous IFN production in A549-SunTag ACE2 screens, modulates detection of CRISPRa-induced ISG effects on SARS-CoV-2. As illustrated in Figure 2D, many of our ISG hits were similarly selected in both *STAT1* genotypes (i.e. both antiviral or both proviral), but failed to clear significance thresholds in one of the screens.

In sum, in conducting focused CRISPRa ISG screens for cell viability effects in A549 cells infected with SARS-CoV-2, we detected several previously known SARS-CoV-2 restriction factors as well as identified new candidate ISGs with putative anti-SARS-CoV-2 activities.

### Validation of screen hits in targeted CRISPRa studies

To confirm hits identified in the pooled screens, we first conducted “single gene” validation experiments with the CRISPRa system for a subset of candidate antiviral ISGs. A549-SunTag ACE2 and A549^ΔSTAT1^-SunTag ACE2 cells were transduced with expression constructs for gRNAs targeting one of eight antiviral ISG hits (*CD74, LY6E, OAS1, CTSS, TRIM25, ERLIN1, ADPRHL2, GBP1*) or with one of two NTGs, treated with Dox to induce gene expression, and infected with SARS-CoV-2 (M.O.I. = 2). Dox-inducible, gRNA-specific CRISPRa-mediated expression for select genes (*CD74, CTSS* and *OAS1*) was confirmed in complementary qRT-PCR experiments (Supplemental figures, 2A-B). We assessed the fraction of infected cells in Dox^On^ cultures (relative to fraction of infected cells in corresponding paired Dox^Off^ cultures, set to 100%) at 24 and 72 hours post-infection. At 24 hours, the fraction of SARS-CoV-2 infected cells (evaluated by flow cytometry for SARS-CoV-2 N protein) was significantly reduced in both A549-SunTag ACE2 and A549^ΔSTAT1^-SunTag ACE2 cultures transduced with gRNAs targeting SARS-CoV-2 restriction factors *CD74*^27^ and *LY6E*^26^ (Fig 3A-B). SARS-CoV-2 infection was also significantly reduced by activation of *OAS1* expression in both A549-SunTag ACE2 and A549^ΔSTAT1^-SunTag ACE2 cultures, while *TRIM25* expression demonstrated significant restriction only in A549-SunTag ACE2 cells (Fig 3A-B). Of note, TRIM25 enhances RIG-I signaling^64^ and has been shown to interact with SARS-CoV-2 RNA^30^, which may suggest that a consequent antiviral effect may require intact IFN signaling. Although some other hits exhibited consistent modest evidence of viral restriction (e.g. *CTSS*), no additional genes met statistical significance thresholds at the 24 hour time point.

**Figure 3:**
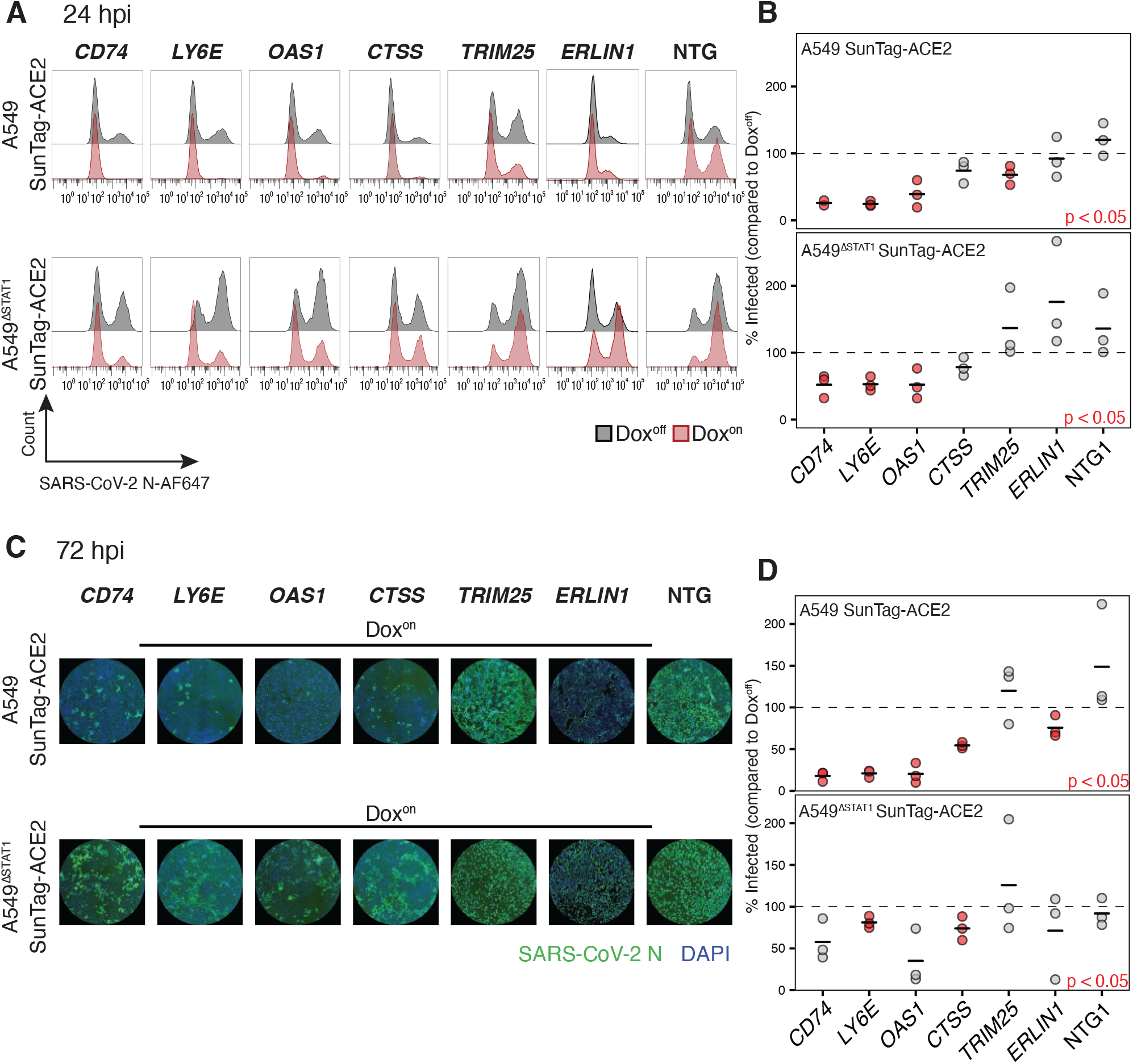
Validation of ISG screen hits in targeted CRISPRa studies. **(A)** Representative flow cytometry histograms for SARS-CoV-2 N protein in A549-SunTag ACE2 and A549^ΔSTAT1^-SunTag ACE2 transduced with indicated gRNAs, treated (red) or not treated (gray) with Dox, at 24 hours post-infection with SARS-CoV-2 (M.O.I. = 2). (**B)** Percent of infected (SARS-CoV-2 N protein positive) cells quantified across biological replicates (n = 3, n = 2 for *CD74* gRNA in A549-SunTag ACE2 cells) for experiments described in **(A)**. Values denote percent of infected cells in Dox^on^ cultures relative to paired Dox^off^ cultures. Points represent individual biological replicates, black lines indicate mean values of biological replicates for each indicated ISG gRNA. Red points indicate statistical significance (p < 0.05) as determined by paired ratio Student’s t-test. **(C)** Representative immunofluorescence images for SARS-CoV-2 N protein and DAPI in A549-SunTag ACE2 and A549^ΔSTAT1^-SunTag ACE2 transduced with indicated gRNAs and treated with Dox, at 72 hours post-infection with SARS-CoV-2 (M.O.I. = 2). **(D)** Percent of infected (SARS-CoV-2 N protein positive) cells quantified across biological replicates (n = 3, n = 2 for ADPRHL2 gRNA in A549-SunTag ACE cells) for experiments described in (C). Values denote percent of infected cells in Dox^on^ cultures relative to paired Dox^off^ cultures. Points represent individual biological replicates, black lines indicate mean values of biological replicates for each indicated ISG gRNA. Statistical significance as in (B).

Several additional hits were confirmed to be antiviral at 72 hours post infection. Once again, the fraction of SARS-CoV-2 infected cells (here assessed by high-throughput microscopy for SARS-CoV-2 N protein due to CPE/fragile cells) was significantly reduced in A549-SunTag ACE2 cultures with activated expression of *CD74, Ly6E*, and *OAS1* (Fig. 3C-D). We also observed significant reduction of infection in cultures expressing *CTSS* in both A549-SunTag ACE2 and A549^ΔSTAT1^-SunTag ACE2 cultures, confirming its activity as a novel SARS-CoV-2 restriction factor. At this time point, *ERLIN1* expression also exhibited modest, yet significant, restriction of SARS-CoV-2 only in A549-SunTag ACE2 cells (Fig. 3C-D). Ectopic expression of *ERLIN1*, a regulator of endoplasmic-reticulum-associated protein degradation (ERAD), was recently shown to restrict SARS-CoV-2 infection^29^. *ADPRHL2* and *GBP1*, additional ISGs identified as antiviral screen hits, did not reach statistical significance in CRISPRa validation experiments (Supplementary Figure 2C-D). Taken together, these results confirm the antiviral effects of multiple ISG screen hits in our CRISPRa system, including *OAS1* and *CTSS*, against SARS-CoV-2.

### Validation of screen hits by ectopic cDNA expression

To further validate antiviral hits confirmed in CRISPRa experiments with a complementary experimental system, we tested ectopically expressed ISG cDNAs for their ability to restrict SARS-CoV-2. A549 ACE2 and A549^ΔSTAT1^ ACE2 cells (both without CRISPRa components) were transduced with Dox-inducible cDNA expression constructs for one of *CD74, LY6E, CTSS, TRIM25, ADPRHL2* or fLuc (Firefly luciferase, negative control; additional independent experiments for *OAS1* cDNAs are described in the following section). After antibiotic selection and expansion, cDNA expression was induced by Dox for 48 hours, after which cells were infected with SARS-CoV-2 (M.O.I. = 2). At 24 hours post-infection, the fraction of infected cells in Dox^On^ and Dox^Off^ conditions was assessed by flow cytometry for SARS-CoV-2 N protein. Dox-inducible cDNA expression of *CD74, LY6E, CTSS* and *TRIM25* recapitulated similar patterns of SARS-CoV-2 restriction (Fig. 4A-B) observed in the CRISPRa system (Fig 3). In addition, *ADPRHL2* also significantly restricted SARS-CoV-2 in cDNA expression experiments (Fig. 4A-B). Interestingly, the apparent antiviral effect conferred by cDNA expression of *CD74* was weaker than that observed in corresponding CRISPRa gRNA-directed expression experiments (Fig. 4A, compared to Fig. 3A). Although these differences could be due to dissimilar expression levels in the CRISPRa and cDNA systems, they might also be explained by isoform-specific effects; the anti-SARS-CoV-2 effects of *CD74* have been assigned to the p41 isoform^27^ that can be expressed in CRISPRa experiments, while the “canonical” isoform p45 (Uniprot identifier P04233-1) was tested in our cDNA experiments. In sum, these experiments offer further validation of the antiviral activity of ISG screen hits against SARS-CoV-2.

**Figure 4:**
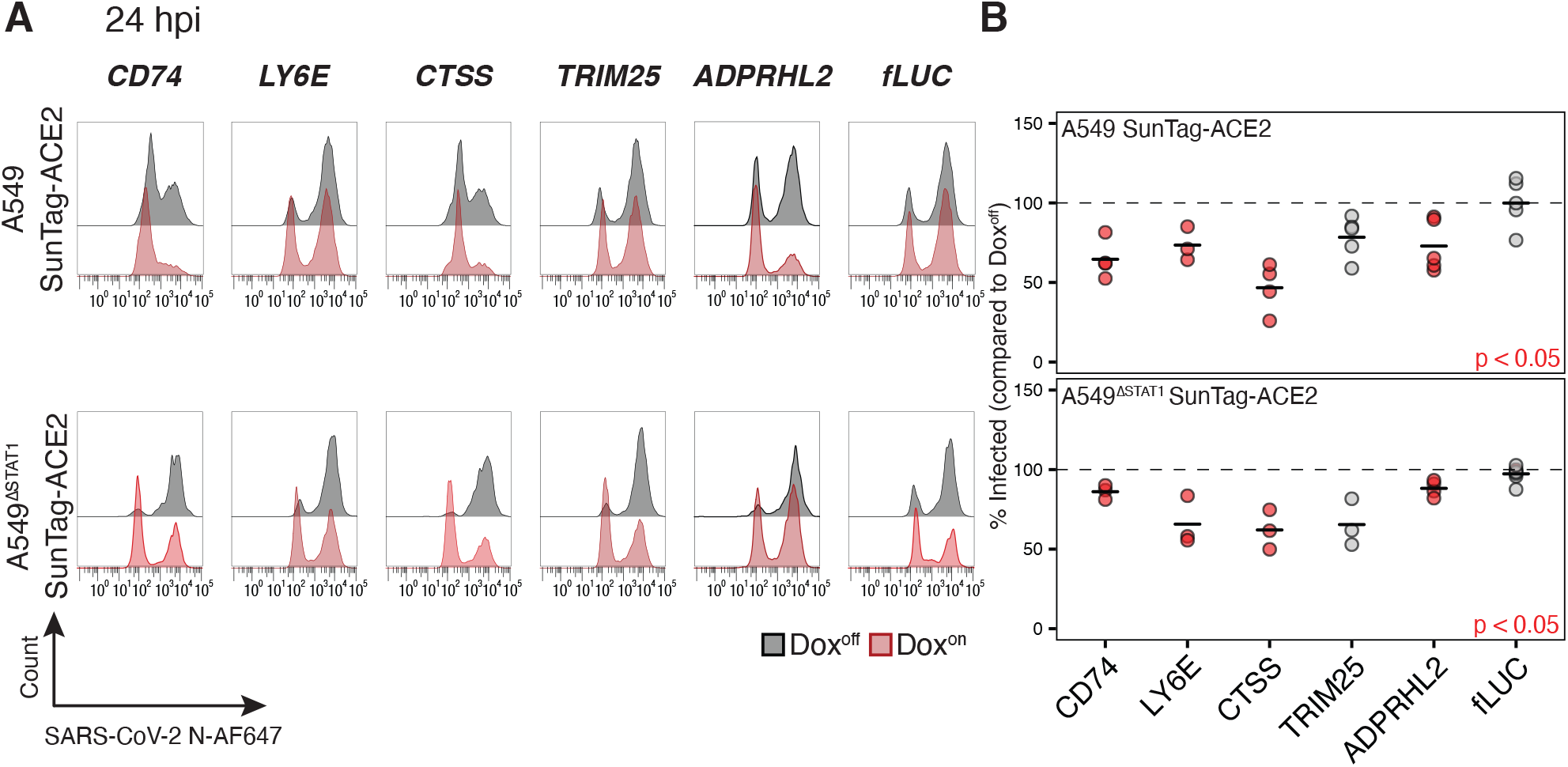
Validation of ISG screen hits by ectopic cDNA expression. **(A)** Representative flow cytometry histograms for SARS-CoV-2 N protein in A549 ACE2 and A549^ΔSTAT1^ ACE2 transduced with expression constructs for indicated cDNAs (fLUC, firefly luciferase negative control), treated (red) or not treated (gray) with Dox, at 24 hours post-infection with SARS-CoV-2 (M.O.I. = 2). **(B)** Percent of infected (SARS-CoV-2 N protein positive) cells quantified across biological replicates (n ≥ 3) for experiments in (A). Values denote percent of infected cells in Dox^on^ cultures relative to paired Dox^off^ cultures. Points represent individual biological replicates, black lines indicate mean values of biological replicates for each indicated ISG cDNA. Red points indicate statistical significance (p < 0.05) as determined by paired ratio Student’s t-test.

### Characterization of OAS1 as a potent SARS-CoV-2 restriction factor

*OAS1* was detected as an antiviral ISG hit in both A549-SunTag and A549^ΔSTAT1^-SunTag screens (Fig. 2) and validated in targeted CRISPRa experiments at both time points tested (Fig. 3). The oligoadenylate synthetase (OAS) family of ISGs are key enzymes involved in antiviral defense^1^. Upon sensing cytosolic dsRNA, OAS proteins are activated to catalyze the formation of 2′-5′-linked oligoadenylate (2′-5′A), which in turn activates latent ribonuclease L (RNaseL). Active RNaseL directly combats diverse viruses by degrading viral genomes, and indirectly by degrading cellular RNA and tRNA^65^. In humans, three members of the OAS family (*OAS1, OAS2, OAS3*) are capable of synthesizing 2′-5′A, and differ by size and sensitivity to dsRNA. An additional family member, *OASL*, is deficient in 2′-5′A catalysis but can sense dsRNA and thereby enhance RIG-I signaling^66,67^. While we detected *OAS1* as a highly ranked hit in both A549-SunTag ACE2 and A549^ΔSTAT1^-SunTag ACE2 screens, other OAS family members were not found to confer antiviral or proviral effects (Fig. 5A).

**Figure 5:**
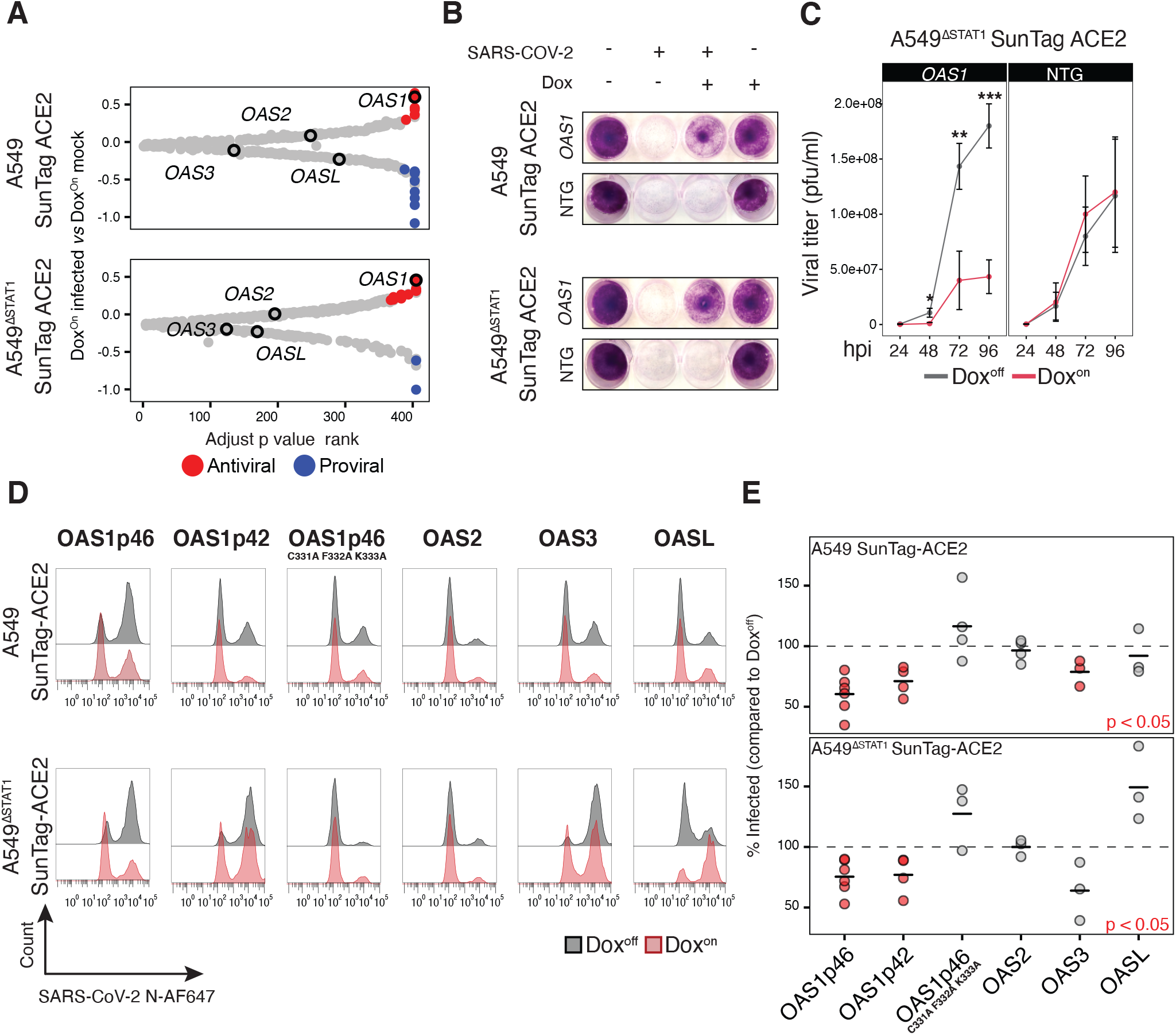
OAS1 is a potent SARS-CoV-2 restriction factor. **(A)** OAS family genes in inducible CRISPRa ISG screen results. For A549-SunTag ACE2 and A549^ΔSTAT1^-SunTag ACE2 screens, ISGs were ranked by SARS-CoV-2:Dox interaction adjusted p value (x axis). β scores for SARS-CoV-2 infection status coefficient (Dox^On^ SARS-CoV-2 infected *vs* Dox^On^ mock infected) are plotted on the y axis, with points passing significance filters are highlighted in red/blue for “antiviral”/“proviral” effects, respectively. OAS family members are labeled as indicated. **(B)** Images of A549-SunTag ACE2 and A549^ΔSTAT1^-SunTag ACE2 cultures transduced with *OAS1* gRNA or NTG, treated/not treated with Dox and infected as indicated with SARS-CoV-2 (M.O.I. = 2, 72 hours infection). Cultures were fixed and stained with Crystal violet to visualize cell viability. **(C)** SARS-CoV-2 growth curves measured by plaque assay. A549^ΔSTAT1^-SunTag ACE2 cells expressing *OAS1* gRNA or NTG gRNA were infected with SARS-CoV-2 (M.O.I. = 0.1) and culture media was collected at indicated time points (x-axis, hours post infection, hpi). Titer was determined by plaque assay on Vero-E6 cells. Statistical significance at each timepoint determined by two-sided Student’s t-test, *** p < 0.0005 ** p < 0.005, * p < 0.05. **(D)** Representative flow cytometry histograms for SARS-CoV-2 N protein in A549 ACE2 and A549^ΔSTAT1^ ACE2 transduced with expression constructs for indicated OAS family member cDNAs, treated (red) or not treated (gray) with Dox, at 24 hours post-infection with SARS-CoV-2 (M.O.I. = 2). **(E)** Percent of infected (SARS-CoV-2 N protein positive) cells quantified across biological replicates for experiments described in **(D)**. Values denote percent of infected cells in Dox^on^ cultures relative to paired Dox^off^ cultures. Points represent individual biological replicates, black lines indicate mean values of biological replicates for each indicated cDNA. Red points indicate statistical significance (p < 0.05) as determined by paired ratio Student’s t-test.

Given its apparent potency in both IFN-responsive and non-responsive contexts, we focused additional studies on characterizing the antiviral activities of OAS1 on SARS-CoV-2 infection. Extending the single gene CRISPRa experimental design described above (Fig. 3), we assessed the effects of gRNA-activated *OAS1* expression on cell viability during SARS-CoV-2 infection. Consistent with results from our positive selection cell survival screens, Dox induction of gRNA-mediated *OAS1* expression conferred a dramatic improvement in cell survival assessed at 72 hours post-infection in both A549-SunTag ACE2 and A549^ΔSTAT1^-SunTag ACE2 cultures; Dox^Off^ cultures and cultures transduced with NTG were readily eliminated by infection (Fig. 5B). In addition to protective effects on cell viability, we also tested the direct impact of *OAS1* expression on the propagation of SARS-CoV-2. To minimize the effects of endogenous IFN production, we infected Dox^Off^ and Dox^On^ A549^ΔSTAT1^ SunTag ACE2 cells expressing *OAS1* or NTG gRNAs and quantified viral titers by plaque assay over time. Activation of *OAS1* expression resulted in a significant decrease in the production of viral progeny, indicating that OAS1 activity functionally restricts SARS-CoV-2 infection (Fig. 5C).

*OAS1* isoforms have been shown to differ in their antiviral activities^36^. To evaluate potential isoform-specific effects of *OAS1* on SARS-CoV-2 restriction, we transduced A549 ACE2 and A549^ΔSTAT1^ ACE2 cells with Dox-inducible expression constructs for cDNAs encoding *OAS1* canonical isoform p46 (*OAS1p46*, Uniprot identifier P00973-1) and a shorter *OAS1* isoform (*OAS1p42*, Uniprot identifier P00973-2) that was recently suggested to be the highest expressed isoform of *OAS1* in A549 cells^36^. To test the requirement of *OAS1* catalytic activity for restricting SARS-CoV-2, we also transduced an expression construct for *OAS1p46* in which amino acids 331, 332 and 333 are replaced with Alanine (*OAS1p46*^C333A/F332A/K333A^). This catalytically inactive mutant is expected to bind dsRNA as well as ATP, but is incapable of the higher order oligomerization required for its enzymatic activity^68^. We also included constructs for *OAS2, OAS3* and *OASL* cDNAs. Following antibiotic selection and expansion, cDNA expression was induced with Dox, cultures were infected with SARS-CoV-2 (M.O.I. = 2) and infection was assessed by flow cytometry (SARS-CoV-2 N protein) at 24 hours post infection. As expected, Dox induction of *OAS1p46* expression significantly reduced the fraction of infected cells in both A549 ACE2 and A549^ΔSTAT1^ ACE2 cells. The p42 isoform of *OAS1* exhibited similar restriction of infection. Inactivation of *OAS1* catalytic activity completely ablated its antiviral effects, suggesting that the mechanism of *OAS1*-mediated SARS-CoV-2 restriction acts through downstream RNaseL activation. Interestingly, while expression of OAS3 exhibited some restriction of SARS-CoV-2 (significant only is A549 ACE2 cells), other OAS family members did not restrict SARS-CoV-2 infection (OAS2), or may have promoted SARS-CoV-2 infection (OASL, only in A549^ΔSTAT1^ ACE2 cells).

### Inducible CRISPRa ISG screen highlights ISGs with antiproliferative/proapoptotic effects

In addition to identification of potential antiviral genes, our experimental design enables indirect assessment of potential ISG effects on cell viability and proliferation outside the context of viral infection. To identify such ISGs, we evaluated significant enrichment/depletion according to their Dox status (Dox^Off^ mock infected *vs* Dox^On^ mock infected, adjusted p value < 0.1, *Supplementary Fig. 3A*). Most significant hits were depleted by Dox treatment (i.e. negatively selected upon expression). Indeed, gRNAs with significantly reduced representation in Dox^on^ conditions included target genes for apoptotic signaling (*TNFRSF10A, TNFAIP3*) and cell cycle negative regulation (*CDKN1A*). The small number of enriched gRNAs (i.e. positively selected upon expression) included Transferrin (*TF*) and Growth differentiation factor 15 (*GDF15*), both of which have been previously described to promote cell proliferation in A549 cells^69,70^. Negatively selected hits common to both A549-SunTag and A549^ΔSTAT1^-SunTag screens included *CDKN1A, TNFRSF10A* and *APOBEC3A*, all previously implicated in reducing proliferation^62,71,72^, and *RNF213, MT1H, UBE2L6, JAK2* and *PXK*, without established roles in cell cycle and/or death pathways (Supplementary Fig. 3B). These results support potential roles for many ISGs, including many with established antiviral activities, in IFN effects on cell viability and proliferation.

## Discussion

Here, we report a CRISPRa strategy for pooled screens of IFN-induced antiviral effectors, and employed this approach to identify ISGs that restrict SARS-CoV-2 cytopathogenicity. Screen results included previously described SARS-CoV-2 restriction factors, as well as multiple additional candidate ISGs with antiviral activity against SARS-CoV-2. Focused CRISPRa and cDNA validation experiments for a subset of these hits confirmed the protective effects of SARS-CoV-2 restriction factors, including *CTSS* and *OAS1*.

### CRISPRa optimized for pooled antiviral ISG screens

CRISPRa strategies have recently been applied to identify genes that regulate viral infection. In a pooled CRISPRa screen, Heaton et al. identified host restriction factors for Influenza A Virus^41^. Using a similar genome-wide gRNA library in a different cell line, Dukhovny et al. detected antiviral effectors with activity against Zika Virus^42^. Recent preprints also reported genome-wide CRISPRa screens for SARS-CoV-2^43,44^. While these examples demonstrate the utility of pooled CRISPRa screens for identifying viral restriction factors genome-wide, we aimed to establish a CRISPRa system optimized for efficiently evaluating ISG activities against respiratory viruses. First, we addressed the potentially confounding effects of endogenous IFN produced by infected cells by expressing the SunTag components in A549^ΔSTAT1^ cells, which are deficient in their capacity to respond to IFN. Indeed, our SARS-CoV-2 CRISPRa ISG screens in A549 and A549^ΔSTAT1^ cells returned almost completely distinct hit lists of candidate antiviral factors; only *OAS1* cleared selection thresholds in both systems. The expression of multiple ISGs can confer additive effects to viral restriction^3^.

Some ISGs suppress cell proliferation or promote apoptosis^1^. These molecular programs may limit viral spread and maintain genome integrity upon detecting nucleic acid damage. Prominent ISGs known to regulate the cell cycle and/or promote cell death include *CDKN1A* (*p21*^71^), *IFI27*^73^, *XAF1*^74^, and members of the oligoadenylate synthetase family of genes^75^. Assessing potential antiviral activities of such genes presents technical challenges, particularly in pooled screen settings in which gene expression alone (and corresponding effects on cell proliferation and/or cell death) is likely to impact relative enrichment/depletion independently of viral infection. Several examples of regulatable CRISPRa systems have been recently described^76–78^. Our implementation of Dox-inducible SunTag CRISPRa gene expression enables transduction and expansion of cultures with gRNAs targeting antiproliferative/proapoptotic ISGs with minimal deleterious effects, as gene expression is only induced shortly before viral infection. Moreover, in comparing Dox^On^ and Dox^Off^ cultures in the absence of viral infection, we are able to assess the antiproliferative/proapoptotic effects of each ISG. Indeed, not only did our results include multiple known cell cycle regulators depleted after only 48 hours after Dox induction, but they also included genes (*CDKN1A, TNFRSF10A* and *OAS1*) with demonstrable effects on *both* virus-independent library enrichment/depletion *and* susceptibility to SARS-CoV-2. While the antiproliferative and proapoptotic effects of IFN are well established^49^, a systematic analysis of the ISGs that mediate these effects remains to be conducted. The initial results and experimental framework described here could be further extended to a comprehensive appraisal of ISG effects on cell cycle and apoptosis. As many cancers exhibit dysregulated ISG expression, such analyses have the potential to inform a variety of therapeutic strategies, particularly oncolytic virus development.

### Cathepsin S restricts SARS-CoV-2

Coronaviruses can enter cells via two different routes: from the cell membrane or from the endosomal compartment. The route of entry is determined in part by the presence of cellular proteases required for spike protein processing^79^. SARS-CoV-2 entry from the cell surface requires *TMPRSS2*, while endosomal entry is mediated by cathepsins that process the spike protein^79^. Cathepsins are cellular proteases that have been implicated in the entry processes of multiple viruses by activating viral glycoproteins to trigger viral fusion at the endosomal membrane^69^. Our screens found that expression of Cathepsin L (*CTSL*), an entry factor for coronaviruses^60^, sensitizes cells to SARS-CoV-2 infection. Intriguingly, Cathepsin S (*CTSS*), an ISG in A549 cells^50^, was identified and validated in our experiments to confer a survival benefit to cells challenged with SARS-CoV-2. Of note, A549 cells lack expression of *TMPRSS2*^80^, making cathepsin glycoprotein processing and endosomal entry likely pathways in viral infection. The apparently opposing effects of *CTSL* and *CTSS* on SARS-CoV-2 infection are surprising and the mechanistic basis for this difference is unclear. *CTSS* and *CTSL* have been shown to bind and cleave different polypeptide motifs^81^, raising the possibility that cleavage by *CTSS* results in suboptimal spike cleavage products that are dysfunctional for viral entry. Alternatively, *CTSS* may interfere indirectly with spike processing by other cathepsins such as *CTSL*. Of note, we found that CTSS maintains its restrictive function in A549^ΔSTAT1^ cells, indicating that its activity against SARS-CoV-2 does not require IFN-induced factors such as *CD74*, which inhibits SARS-CoV-2 by blocking cathepsin-mediated entry^27^.

### OAS1 is a potent SARS-CoV-2 restriction factor

Oligoadenylate synthetase family members are broadly acting ISGs important for innate antiviral defense against multiple viruses^66^. RNaseL, the downstream effector of *OAS1-3*, degrades cellular and viral RNA upon activation and thereby limits viral propagation. RNaseL activity has been directly implicated in host defense against different coronaviruses^82,83^, most recently against SARS-CoV-2^84^. The OAS/RNaseL pathway is antagonized by MERS-CoV, which blocks RNaseL activation by degrading 2′-5′A species generated by OAS proteins^82^. Our screens identified *OAS1* as a SARS-CoV-2 restriction factor in both A549-SunTag ACE2 and A549^ΔSTAT1^-SunTag ACE2 cells. Our experiments further demonstrated that *OAS1* catalytic activity is necessary for its effects on SARS-CoV-2. This observation suggests that SARS-CoV-2 may not directly antagonize the generation of 2′-5′A like MERS-CoV, and therefore remains susceptible to RNaseL effector functions^84^.

Although *OAS1* has not been identified as a SARS-CoV-2 restriction factor in other recent SARS-CoV-2 ISG screens^29,85,86^, it was recently shown that ectopic *OAS1* expression in 293T cells can effectively interfere with SARS-CoV-2 propagation^87^. Moreover, a growing body of genetic, epidemiological, and clinical data support an important role for *OAS1* in both SARS-CoV and SARS-CoV-2 host defense. *OAS1* genetic variants were linked to infection and excessive morbidity in the SARS-CoV outbreak^88,89^. More recently, genome-wide association studies (GWAS) have associated single nucleotide polymorphisms (SNPs) in *OAS* loci with COVID-19 mortality^90^. Clinical studies have shown that elevated levels of plasma *OAS1* are associated with reduced COVID-19 hospitalization and mortality; these effects are amplified by an *OAS1* isoform of Neanderthal origins^91^.

Given the apparently potent effects of *OAS1* in our experiments, we were surprised that it had not been detected in other recently published genome-wide and ISG-focused screens^29,43,44,85,86^. This discrepancy could be due to the distinct features of the experimental systems used, including different cell types, different infection time points, and low M.O.I. infections of IFN-competent cell lines, in which potential paracrine signaling may obscure certain ISG effects. This possibility is generally supported by the relatively few ISGs detected in pooled activation screens^41–44^, which would otherwise be predicted to be enriched due to their direct antiviral effects. Moreover, as *OAS1* has been characterized as proapoptotic^92^, our inducible system and multifactorial analysis strategy may have been uniquely capable to robustly detect its effects.

**Supplemental figure S1:**
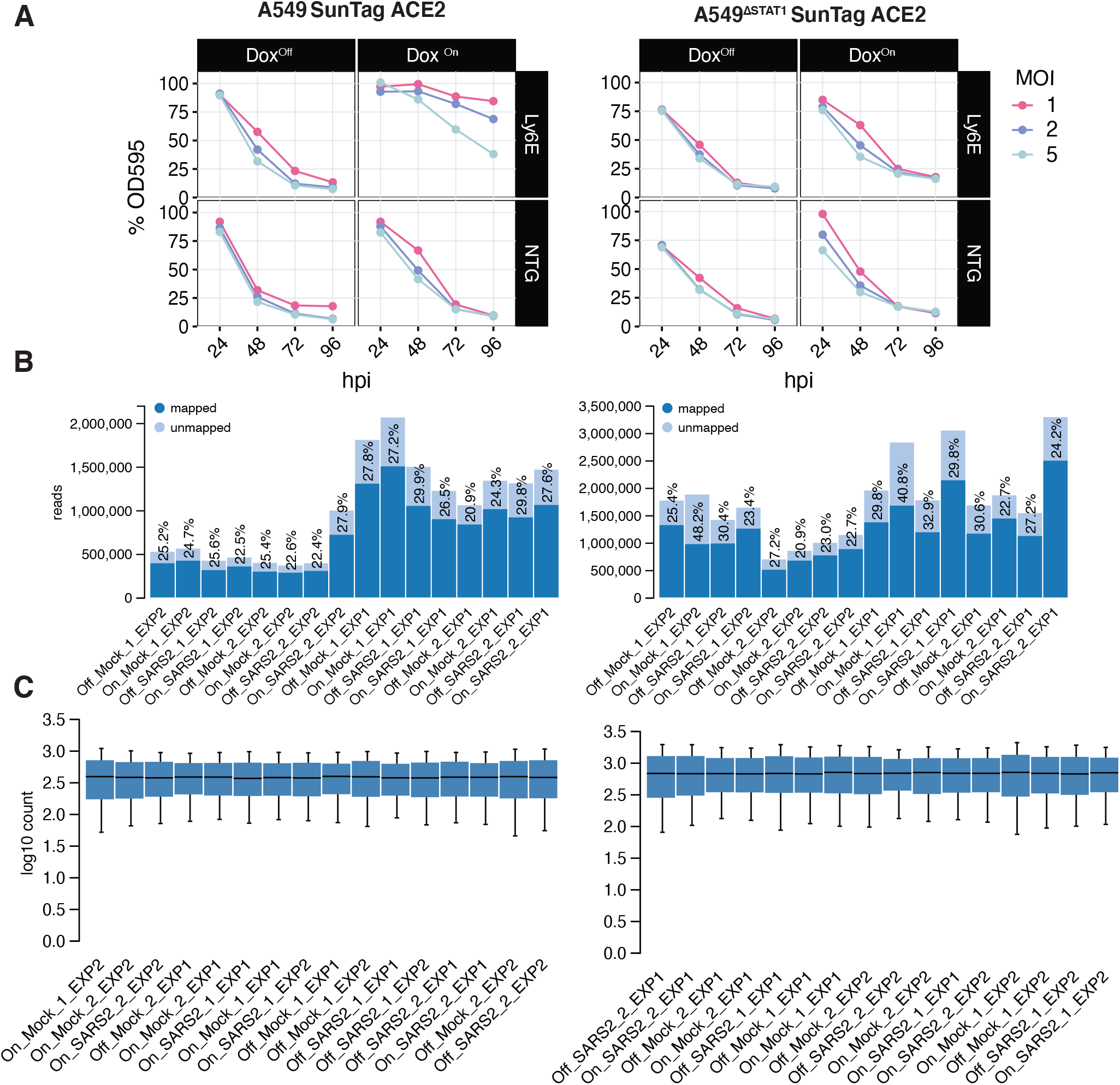
Inducible CRISPRa ISG screen optimization and quality control. **(A)** Pilot experiment using Methylene Blue assay to assess SARS-CoV-2 CPE under different infection conditions. A549-SunTag ACE2 and A549^ΔSTAT1^-SunTag ACE2 cells, expressing *LY6E* gRNA or a non-targeting gRNA (NTG) were infected with SARS-CoV-2 at indicated M.O.I., fixed at indicated time points, and stained with methylene blue. Values indicate percent OD595 absorption relative to time point = 0 (set to 100%). **(B)** CRISPRa screen quality metrics: sequencing reads per sample. Values indicate number of reads sequenced for each sample in the pooled screens. Percentage values (light fill) for reads that fail to map to gRNA sequences in the ISG library reference. **(C)** CRISPRa screen quality metrics: Normalized read count distribution per sample. Log_10_ transformed read count for each sample normalized to the count of non-targeting guides.

**Supplemental figure S2:**
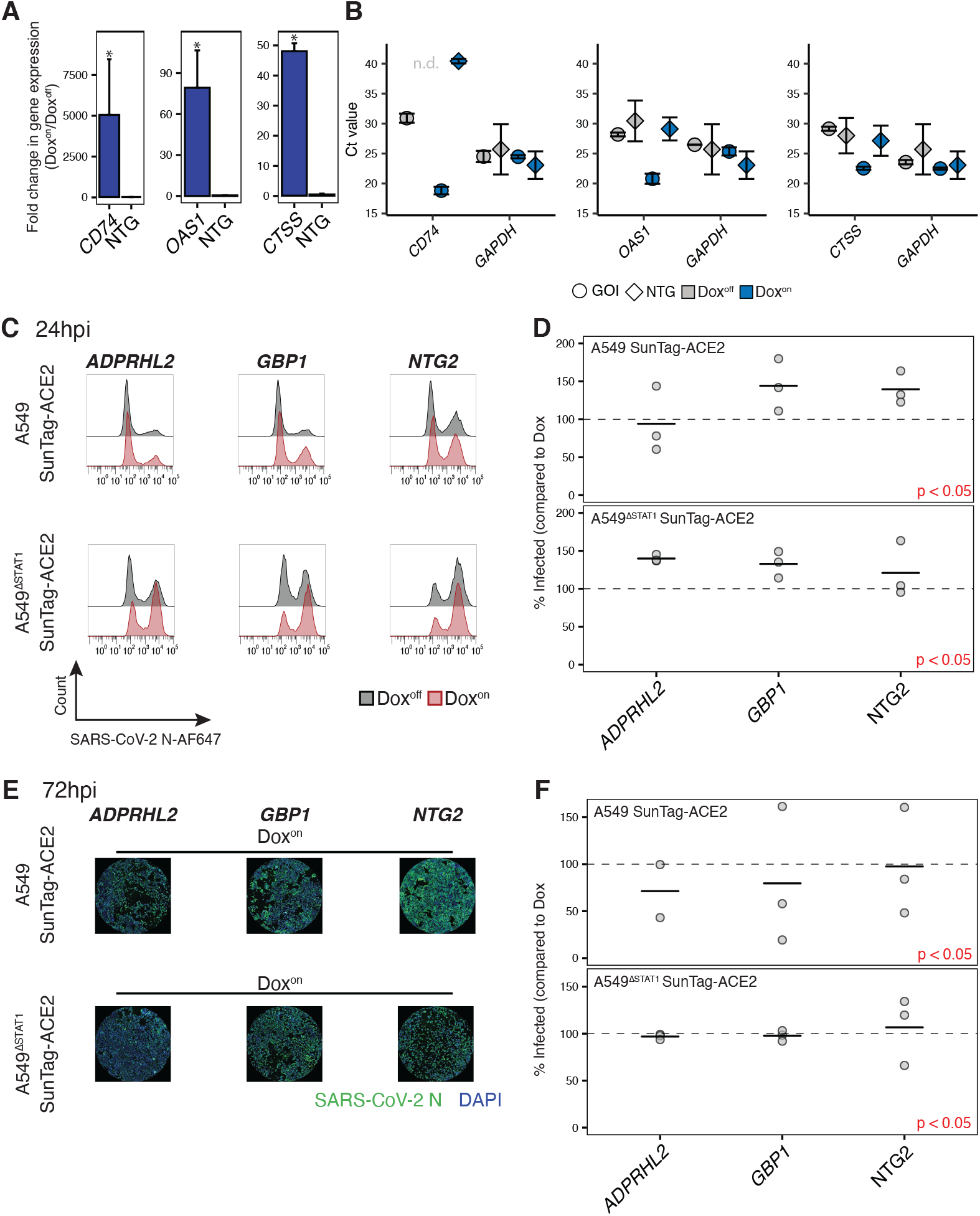
Complete validation results of screen hits tested in targeted CRISPRa studies. **(A)** qRT-PCR analysis demonstrates effective CRISPRa gene induction in A549-SunTag ACE2 cells. Values indicate fold change expression (Dox^on^ relative to Dox^off^) for indicated genes (*CD74, OAS1* and *CTSS*) in A549-SunTag ACE2 cells expressing either corresponding gRNA or NTG. Fold change values calculated using the ΔΔCt method with GAPDH as normalization control. Undetectable Ct value for Dox^off^ condition of cells expressing NTG and probed for *CD74* expression was set to 40 to enable fold change calculation. **(B)** qRT-PCR mean threshold cycle (Ct) values for *CD74, OAS1, CTSS* and *GAPDH* in A549-SunTag ACE2 cells expressing gRNA against gene of interest (GOI, circle) or NTG (diamond) in Dox^on^ and Dox^off^ cells. Error bars indicate ± SD Ct value. **(C)** Representative flow cytometry histograms for SARS-CoV-2 N protein in A549-SunTag ACE2 and A549^ΔSTAT1^-SunTag ACE2 transduced with indicated gRNAs, treated (red) or not treated (gray) with Dox, at 24 hours post-infection with SARS-CoV-2 (M.O.I. = 2). **(D)** Percent of infected (SARS-CoV-2 N protein positive) cells quantified across biological replicates (n = 3) for experiments described in **(C)**. Values denote percent of infected cells in Dox^on^ cultures relative to paired Dox^off^ cultures. Points represent individual biological replicates, black lines indicate mean values of biological replicates for each indicated ISG gRNA. Red points indicate statistical significance (p < 0.05) as determined by paired ratio Student’s t-test. **(E)** Representative immunofluorescence images for SARS-CoV-2 N protein and DAPI in A549-SunTag ACE2 and A549^ΔSTAT1^-SunTag ACE2 transduced with indicated gRNAs and treated with Dox, at 72 hours post-infection with SARS-CoV-2 (M.O.I. = 2). **(F)** Percent of infected (SARS-CoV-2 N protein positive) cells quantified across biological replicates for experiments described in (E). Values denote percent of infected cells in Dox^on^ cultures relative to paired Dox^off^ cultures. Points represent individual biological replicates, black lines indicate mean values of biological replicates for each indicated ISG gRNA. Statistical significance as in (D).

**Supplemental figure S3:**
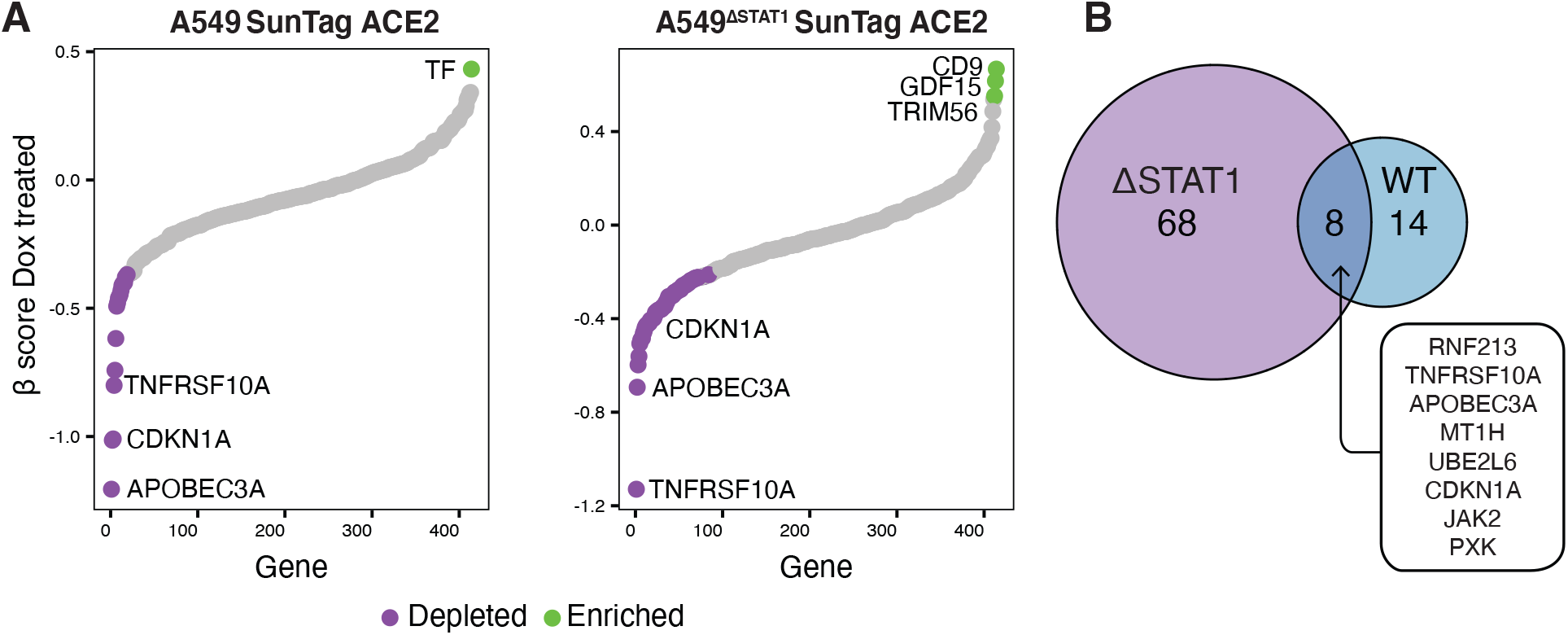
Inducible CRISPRa ISG screen highlights ISGs with putative antiproliferative/proapoptotic effects. **(A)** ISGs ranked by the inverse of their β-scores for the Dox status coefficient (Dox^Off^ mock infected *vs* Dox^On^ mock infected). Significantly (adjusted p value < 0.1) enriched/depleted gRNAs are highlighted in green/purple respectively. **(B)** Venn diagram of significantly depleted (i.e., candidate antiproliferative/proapoptotic) ISG hits from (A) in A549-SunTag ACE2 and A549^ΔSTAT1^-SunTag ACE2 screens.

**Supplemental figure S4:**
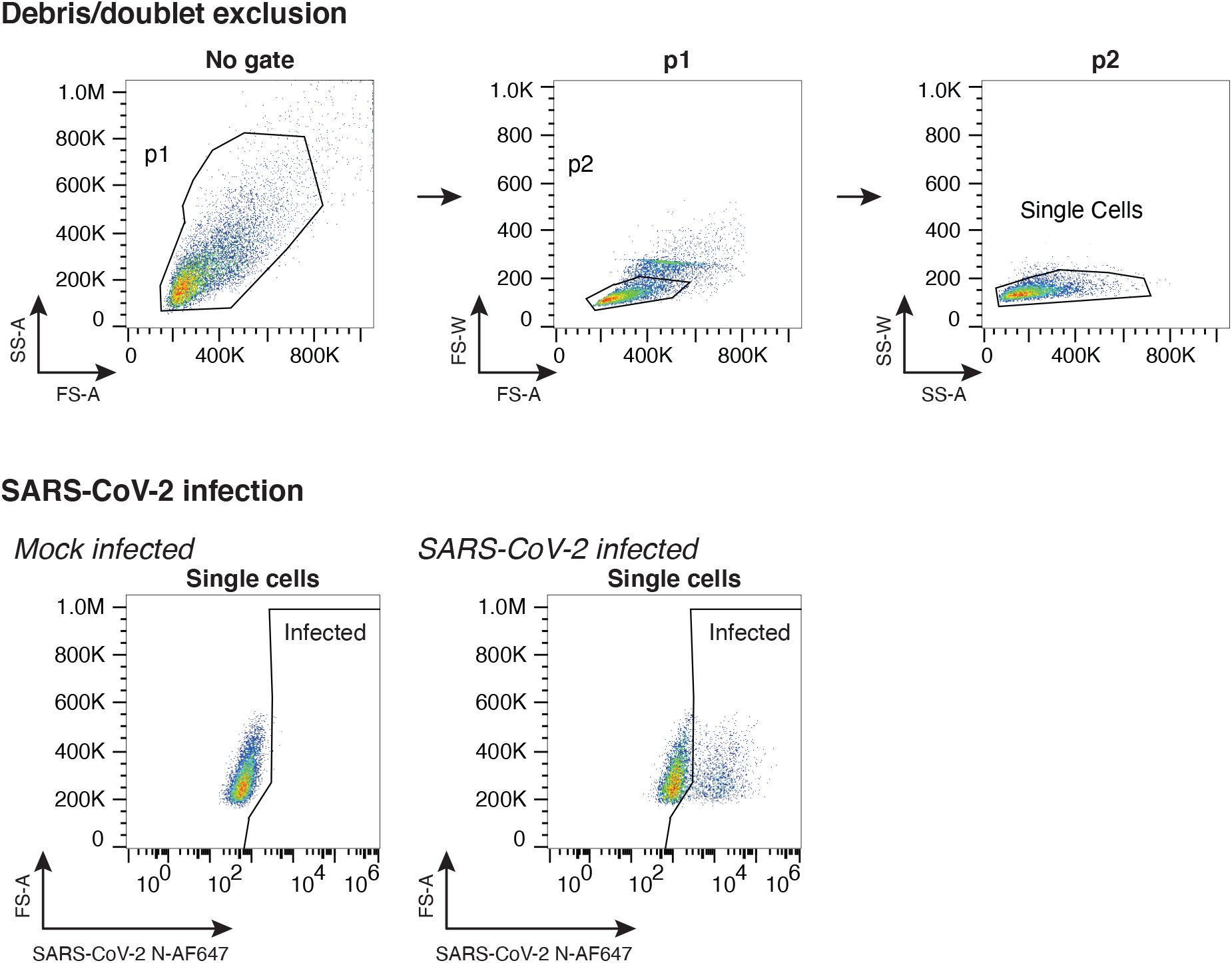
Gating strategy for flow cytometry experiments. Representative gating strategy for identifying SARS-CoV-2 infected cells.

## Materials and methods

### Cell lines and VSV-GFP

All cell lines used in this study were maintained in Dulbecco’s Modified Eagle Medium (DMEM, Corning #10-017-CV) supplemented with 10% fetal-bovine serum (FBS) and 1% Penicillin Streptomycin (PSN, Fisher scientific #15-140-122), and routinely cultured at 37° C with 5% CO_2_. Vero-E6 cells (ATCC, CRL-1586) were used for propagation of SARS-COV-2 and for plaque-assays. Lenti-X 293T cells (Takara #632180) were used for lentivirus packaging. A549 and A549^ΔSTAT1 50^, a kind gift from Dr. Meike Dittmann (NYU Langone School of Medicine), were used for generation of CRISPRa cell lines and for infection studies. A549 and derived cell lines (ACE2-expressing cell lines with or without CRISPRa expression and ΔSTAT1^50^ counterparts) were validated by short tandem repeat (STR) analysis (all confirmed 100% match to A549, CVCL_0023). All cell lines were routinely tested (Boca Scientific #50-168-5641) negative for Mycoplasma contamination. Recombinant-Indiana vesiculovirus expressing GFP (VSV-GFP) was a gift from Dr. Dusan Bogunovic (Icahn School of Medicine at Mount Sinai). Propagation and infections were conducted as previously described^52^.

### Reagents and chemicals

For doxycycline treatments to induce gene expression (both CRISPRa cell lines and cDNA ORF expression constructs), cultures were incubated in DMEM 10% FBS supplemented with 5μg/ml Doxycycline (Sigma # 50-165-6938) for 48 hours prior to infection or culture harvest as indicated for each experiment. Infections (VSV-GFP, SARS-CoV-2) were performed for 1 hour at 37°C in the absence of Dox, which was re-added (1μg/ml) upon removal of the virus inoculum. For IFN experiments, cultures were treated with 200U of recombinant human IFNα2b (PBL Assay Science # 11105-1) for 3 hours prior to culture harvest.

### Propagation and titration of SARS-COV-2

SARS-COV-2 (isolate USA-WA1/2020, BEI resource NR-52281) stocks were grown by inoculating confluent T175 flasks of Vero-E6 cells with SARS-COV-2 isolate (passage 2). Infected cultures were maintained in reduced-serum DMEM (2% FBS) for 3 days, after which medium was collected and filtered by centrifugation (8000 x g, 15 minutes) using an Amicon Ultra-15 filter unit with a 100KDa cutoff filter (Millipore # UFC910024). Concentrated virus stocks in reduced-serum DMEM (2% FBS) supplemented with 50mM HEPES buffer (Gibco #15630080) were stored at -80°C. To determine the number of infectious units in each viral stock (IU/ml), target cell lines (A549-ACE2 or A549-SunTag-ACE2) were plated in duplicate in 24 well plates, and were infected with 2-fold serial dilution series of SARS-CoV-2 stocks at 37°C for 1 hour, after which virus was removed and replaced with DMEM supplemented with 10% FBS. At 24 hours post-infection, cultures were harvested and fixed by incubating in 4% Paraformaldehyde (Alfa Aesar #AA433689M) in PBS for 24 hours. The fraction of infected cells was determined by flow cytometry for SARS-CoV-2 N protein (details below). The percentage of infected cells was used to determine the IU/ml values for each viral stock by using the formula:

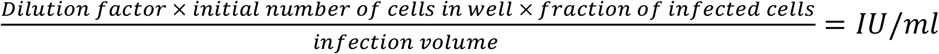

All SARS-CoV-2 propagations and experiments were performed in a biosafety level 3 facility in compliance with institutional protocols and federal guidelines.

### Generation of lentiviruses and viral transduction

To generate lentiviruses for gRNA or cDNA expression, a mix of 2.5μg of the desired transfer vector, 2μg psPAX2 and 0.8μg of pMD2.G (a gift from Didier Trono, Addgene #12260 and #12259, respectively), 14μl Lipofectamine 3000 (Fisher scientific #L3000001) and 20μl of P3000 reagent, was prepared in 250μl of OptiMEM (Gibco #11058021). This transfection mix was added to 1.5×10^6^ Lenti-X 293T cells (Takara Bio #632180, plated in 6 well plates 18 hours prior to transfection) in1 ml of 10% FBS DMEM for 8 hours after which transfection media was removed and replaced with 2ml of 10% FBS DMEM. 3 days post-transfection lentivirus supernatants were collected and centrifuged at 800g for 5 minutes to pellet cell debris, filtered through 45μm PVDF filters (Millipore # SLHVR33RS), and stored at -80°C. Transductions were performed on semi-confluent wells in the presence of Polybrene (8μg/ml, Signa #H9268) using spin-infection (800 x g, 37°C, 90 minutes). Culture media containing selection antibiotics was added to transduced cultures 24 hours post-transduction.

### Cloning of inducible CRISPRa-SunTag system components

To generate the plasmids for the inducible CRISPRa system, we re-engineered an existing CRISPRa technology^45^ to enable Dox-inducible expression and independent component construct antibiotic selection. pCW-TRE, a Dox-inducible expression vector, was generated by modifying pCW-Cas9-Blast (a gift from Mohan Babu, Addgene # 83481) to include a single BamHi site. Next, the antibody component of the SunTag system (pHRdSV40-scFv-GCN4-sfGFP-VP64-GB1-NLS^45^, a gift from Ron Vale, Addgene #60904) was digested with EcoRI+NotI and subcloned into the blunted BamHi site of pCW-TRE. The nuclease-inactive Cas9 fused to the SunTag scaffold (dCas9-SunTag derived from pHRdSV40-dCas9-10xGCN4_v4-P2A-BFP^45^, a gift from Ron Vale, Addgene #60903) was assembled in-frame with a hygromycin resistance gene into pHR-PGK (a gift from Wendell Lim, Addgene #79120^93^), generating pHR-PGK-dCas9-SunTag-P2A-HygR.

### Generation of A549-SunTag and A549^ΔSTAT1^-SunTag cells

To generate A549-SunTag and A549^ΔSTAT1^-SunTag cells, we transduced A549 or A549^ΔSTAT1^ cells with lentiviruses encoding the Dox-inducible transactivator component of SunTag (pCW-TRE-scFv-SunTag). Single cell clones were selected with Blasticidin (Fisher Scientific #BP264725, 1μg/ml). Next, pCW-TRE-scFv-SunTag single cell clones were transduced with lentiviruses encoding the SunTag nuclease-inactive Cas9 (pHR-PGK-dCas9-SunTag-P2A-HygR) and single cell clones were selected with Hygromycin (Thermo Scientific 10687010, 500μg/ml). Finally, multiple single cell clones of A549-SunTag and A549^ΔSTAT1^-SunTag, selected for the expression of both components, were tested for gRNA-directed gene activation capabilities with gRNAs targeting *CXCR4* and measuring *CXCR4* protein expression by flow cytometry as previously described^45^.

### Generation of ACE2 expressing cell lines

To generate ACE2-expressing A549 cell lines, the human ACE2 coding sequence (RefSeq accession NM_001371415.1) was PCR amplified and cloned into the BamHi site of lentiviral vector pHR-PGK (Addgene #79120). Lentivirus was produced as described above and used to transduce 5×10^4^ target cells (A549, A549^ΔSTAT1^, A549-SunTag or A549^ΔSTAT1^-SunTag) in 12 well plates. Single cell clones were expanded and validated for expression of ACE2 by Western Blot analysis (Abcam #ab15348).

### qRT-PCR

For indicated experiments, A549 cultures were harvested by trypsinization, pelleted, and homogenized in Trizol (Invitrogen #15596026). Total RNA was isolated with the Direct-zol RNA MiniPrep kit (Zymo #R2050), including DNase protocol, according to manufacturer’s instructions. For each sample, 1μg of total RNA was reverse transcribed (Applied Biosystems #4368814) with random hexamers for priming. qRT-PCR was performed with the TaqMan Universal Master Mix (Applied Biosystems #4440038), and TaqMan primer/probe sets for each gene of interest (*MX1*; Hs00895608_m1, *CD74*; Hs00269961_m1, *OAS1*; Hs00242943_m1, *CTSS*; Hs00175407_m1, *GAPDH*; Hs99999905_m1). Ct values were used for calculation of gene expression using the comparative threshold cycle method^94^ with *GAPDH* expression as loading control, comparing Dox^on^ condition to Dox^off^ condition.

### SARS-CoV-2 infections

For validation studies employing gRNA or cDNA to express ISGs, SARS-CoV-2 stocks were diluted in reduced serum DMEM (2% FBS) supplemented with 50mM HEPES and 1% PSN, inoculated to indicated cell cultures, and incubated at 37°C for 1 hour. Infection medium was then replaced with DMEM (10% FBS and 1% PSN) for timepoints indicated in each experiment.

### Methylene blue assay

A549-SunTag and A549^ΔSTAT1^-SunTag cells were plated in 96-well plates (3000 cells/well, 4 replicates per condition, 1 plate for each time point) and infected with SARS-CoV-2. Infections were done in reduced serum DMEM (2% FBS, 50mM HEPES, 1% PSN), and virus was left on the cells for the indicated time points. Plates were fixed in 4% Paraformaldehyde at room temperature for a minimum of 24 hours. Cells were then washed twice with 100μl 0.1M sodium tetraborate (Sigma # 221732), stained with 0.5% methylene blue (Sigma # M9140) in 0.1M sodium tetraborate (15 minutes, room temperature), extensively washed in 0.1M sodium tetraborate, and extracted with 0.1M HCl. Absorbance was measured on a BioTek Cytation plate reader at 595 nm.

### Curation and Cloning of the ISG library

To assemble the list of ISGs targeted by the gRNA library, an established list of ISGs^3,4^ was combined with a list of genes upregulated (RNA-Seq log_2_ fold-change >2, adjusted p value < 0.05) after 6 or 48 hours of IFN stimulation in A549 cells^50^. To focus the list on direct antiviral effectors, genes annotated as transcription factors^55^, HLA genes, and central PRRs were excluded. gRNA sequences (n = 3 per gene) for the resulting 414 gene list were selected from the Calabrese library^56^. The final gRNA library pool contained 1242 ISG-targeting gRNAs and 24 non-targeting controls. The gRNA library was synthesized as an oligonucleotide pool (Integrated DNA Technologies) and cloned into CROP-seq-opti (a gift from Jay Shendure, Addgene # 106280^95^) as described^96^. Briefly, the CROP-seq-opti backbone was digested with BsmBi and gel purified. 2000 fmoles of the gRNA library oligonucleotide pool were mixed with 50 fmoles of linearized CROP-seq-opti in 10μl of NEB-Builder master mix (New England Biolabs # E2621). After 1 hour incubation at 50°C, the assembled plasmid pool was used to transform 25μl of electrocompetent bacteria (Lucigen #60242-2) on a Bio-Rad Gene-Pulser 2 electroporation system (Bio-Rad # 1652105) with the following settings: 25 μF, 200 Ohm, 1.5KV. Ampicillin resistant colonies were pooled, grown overnight in liquid culture (LB broth, Fisher BioReagents #BP1426) at 32°C, and the plasmid library was extracted by Maxi prep (Qiagen #12362) according to the manufacturer’s protocol. Plasmid library was packaged into lentiviruses and transducing units/ml (TU/ml) were determined by calculating colony forming units/ml of Puromycin (Sigma-Aldrich #P8833, 2μg/ml)-resistant transduced A549 cultures.

### Inducible CRISPRa ISG screen for SARS-CoV-2 restriction factors

For each of 2 clones from each *STAT1* genotype (A549-SunTag ACE2 or A549^ΔSTAT1^-SunTag ACE2, 2 clones each), 6 × 10^6^ cells were transduced with gRNA library lentivirus (described above) at M.O.I. = 0.1, assuring zero or one gRNA/cell in more than 95% of library cells, and representation of 500 cells/gRNA. Transduced cultures were selected for CROP-seq-opti transduction with Puromycin (Sigma-Aldrich #P8833, 2μg/ml), and expanded for 14 days. 48 hours prior to SARS-COV-2 infection, gene expression was induced by treating the cells with Dox (5μg/ml). 2 × 10^6^ cells (estimated representation of more that 1500 cells/gRNA) were infected with SARS-COV-2 (M.O.I. = 3). 72 hours post infection, after observation of significant CPE in infected cultures, genomic DNA was extracted from surviving cells for Illumina sequencing library preparation. In total, screens were conducted in two experiments, each experiment including two single cell clones from each *STAT1* genotype. In the first experiment, following infection, an additional 4ml of reduced-serum DMEM were added to the cultures, but the virus inoculum was not removed from culture wells. In the second experiment, the virus inoculum was removed from the cells after 1 hour of infection, cultures were washed twice with calcium/magnesium-free PBS, and cultured in DMEM (10% FBS) for 72 hours.

### Screen library preparation and sequencing

CROP-seq-opti gRNA sequencing libraries were prepared as previously described^96^. Briefly, 100ng of each gDNA sample was PCR amplified in triplicate with Q5 High-Fidelity DNA Polymerase (NEB #M0494S) with 500nM primers (Supplemental table 4) flanking the guide sequence cassette and including Illumina adaptor sequences and sample index sequences (98°C x 30s, 98°C x 10s, 72°C x 45s, 25 cycles). PCR products were purified using 2.0X AMPure XP magnetic beads (Beckman Coulter #A63881) according to manufacturer’s protocol. Sequencing libraries were quantified with the KAPA Library Quantification Kit (Roche #07960140001), pooled, and sequenced in multiplex on the Illumina NextSeq 500 platform using a 150-cycle mid output kit (Illumina # 20024904) with read configuration of 167 bases (read 1) and 8 bases (i7 index).

### Inducible CRISPRa ISG screen data processing and analysis

Illumina BCL sequence files were converted to FASTQ format with the bcl2fastq tool (v2.20.0.422, Illumina). gRNA enrichment/depletion analyses were conducted with the MAGeCK package (version 0.5.9.4^57^) Sample x gRNA count matrixes were generated with the MAGeCK *count* function. Enrichment/depletion analyses were conducted with MAGeCK MLE for each *STAT1* genotype. To analyze gRNA enrichment, we used MAGeCK MLE with a design matrix generated by the R (version 4.0.2) *model*.*matrix()* function, with design formula specified as: ∼(dox*virus) + experiment + clone. The resulting generalized linear model included factors for Dox status, SARS-CoV-2 infection status, clone, and experiment, as well as a dox:virus interaction term (used to test for gRNA enrichment/depletion by virus infection differences by dox status). MAGeCK MLE was run with 10 permutations, with normalization to NTG gRNAs. Enrichment/depletion p values were adjusted by the method of Benjamini and Hochberg within the MAGeCK MLE framework. To focus on genes enriched or depleted upon viral infection beyond potential virus-independent effects on library representation, we filtered to include hits with adjusted p value less than 0.1 for both SARS-CoV-2 infection status coefficient (Dox^On^ SARS-CoV-2 infected *vs* Dox^On^ mock infected) and the Dox status:SARS-CoV-2 infection status interaction term. “Antiviral”/ “proviral” designations were made based on the sign of the β score for the SARS-CoV-2 infection status coefficient.

In analyses for antiproliferative/proapoptotic ISGs, hits were selected as adjusted p value less than 0.1 for the Dox status coefficient (Dox^Off^ mock infected *vs* Dox^On^ mock infected), and β score were multiplied by -1 to facilitate enrichment/depletion interpretation within the required model syntax.

### Cloning procedures for individual guide RNAs

Cloning of individual gRNAs into CROP-seq-opti vectors was performed as previously described^96,97^. Briefly, for targeted (i.e. non-library) CRISPRa experiments, gRNA sequences were synthesized (Integrated DNA technologies) as oligonucleotide duplexes with BsmBI-compatible overhangs. CROP-seq-opti (Addgene # 106280) was linearized by BsmBI (New England Biolabs # R0580S) digestion. Oligonucleotides were phosphorylated with T4 polynucleotide kinase (New England Biolabs #M0201L), annealed and ligated (Quick Ligation kit, New England Biolabs #M2200S) into BsmBI digested CROP-seq-opti. 2μl from the ligation reaction were used for transformation of 10μl of NEB stable competent E. coli cells (C3040, New England Biolabs # C3040H). Proper insertion of gRNA sequence was confirmed by Sanger sequencing primed from the U6 promoter region.

### Cloning procedures for ISG cDNA ORFs

To clone screen hit cDNAs for validation experiments, the complete coding sequences of canonical isoforms (annotated by Uniprot; Bateman et al., 2021) of candidate genes were either synthesized as gBlocks (Integrated DNA Technologies) or PCR amplified from cDNA derived from IFNα2b-treated A549 cells. Genetic sequences were synthesized/amplified with homology overhangs complementary to the overhangs of EcoRI+BamHi digested pLVX-TetOne-Puro (Takara # 631849). 100ng of cDNA were ligated into 75ng of gel-purified digested vector using Neb builder (New England Biolabs #E2621) in a final reaction volume of 10μl. Ampicillin resistant colonies were grown overnight in LB media at 30°, and plasmids were extracted by Mini prep (Qiagen # 27106) according to the manufacturer’s protocol.

### Flow cytometry

For VSV-GFP experiments, cells were fixed with 4% Paraformaldehyde for 30 minutes at room temperature and analyzed for GFP fluorescence on a Gallios flow cytometer (Beckman-Coulter). For SARS-CoV-2 experiments, cells were fixed with 4% paraformaldehyde at room temperature for a minimum of 24 hours, washed once with PBS and permeabilized with 1X perm-wash buffer (BDBiosciences #554723) for 5 minutes. AlexaFluor 647-conjugated SARS-CoV nucleocapsid (N) antibody (clone 1C7C7, generously provided by the Center for Therapeutic Antibody Discovery at the Icahn School of Medicine at Mount Sinai) was diluted 1:400 in perm-wash buffer, and added directly to permeabilized samples, which were then incubated at room temperature for 40 minutes in the dark. Samples were washed once with 1X perm-wash buffer, once with calcium/magnesium-free PBS, and acquired on a Gallios flow cytometer (Beckman-Coulter). For all viral infections, analysis was performed with FlowJo software (Version 10.7.1, Becton Dickinson), excluding cell doublets and debris and gating according to mock infected populations (Supplementary Fig. 4). Samples with fewer than 2000 cell events after doublet and debris gating were excluded from analysis.

### Immunofluorescence high throughput microscopy

SARS-CoV-2 infected A549-SunTag ACE2 and A549^ΔSTAT1^-SunTag ACE2 in 96-well plates were fixed with 4% Paraformaldehyde at room temperature for a minimum 24 hours, washed with PBS, and permeabilized with 0.1% Triton X-100 (Fisher scientific # AC327371000) for 15 minutes. Plates were blocked with 3% Bovine serum albumin (BSA, Miltenyi Biotec #130-091-376) in PBS for 1 hour. AlexaFluor 647-conjugated SARS-CoV nucleocapsid (N) antibody (clone 1C7C7) was diluted 1:2000 in 0.5% BSA in PBS and added to wells for 1 hour incubation. Samples were then washed twice with PBS and stained with 4′,6-diamidino-2-phenylindole (DAPI, Thermo Scientific #D1306) diluted 1:200 in 0.5% BSA in PBS. All steps were done at room temperature. Plates were imaged on a Celigo instrument (Nexcelom Biosciences) and the fraction of infected cells per well was determined using Cell Profiler^99^.

### SARS-CoV-2 growth curves by plaque assay

A549^1STAT1^ SunTag cells, expressing gRNAs targeting either *OAS1* or NTG, were pretreated, or not, with Dox (5μg/ml, 48 hours before infection), and infected with SARS-CoV-2 (M.O.I. = 0.1). After 1 hour, virus inoculum was removed and replaced with DMEM/10% FBS supplemented with or without Dox. Every 24 hours post infection, 100μl of the culture supernatants was collected, serially diluted, and used to infect 2×10^5^ Vero-E6 cells plated in a 24 well plates (37°C, agitating every 10 minutes). After 1 hour, inoculum was removed and replaced with overlay media consisting of Minimum Essential Media (MEM, Thermo # 11095080) supplemented with 0.8% SeaPlaque Agarose (Lonza # 50104), 4% FBS and 1% PSN pre-warmed to 37°. At 72 hours post-infection, Vero-E6 cells were fixed in 4% Paraformaldehyde at room temperature for 24 hours, washed twice with PBS and stained with 1% Crystal violet (Sigma # C0775) for 15 minutes. Viral titer was determined by calculating infectious units/ml.

### Quantification and statistical analysis

Unless otherwise indicated, error bars indicate standard deviation from the mean (SD) of at least 3 biological replicates. For flow cytometry and immunofluorescence experiments, statistical significance was determined with one-sided paired ratio Student’s t-test, with p values < 0.05 considered to be significant.

## Supporting information

Supplemental Table 1 List of ISGs

Supplemental Table 2 ISG library oligo pool

Supplemental Table 3 Screen results MAGEeCK-MLE output

Supplemental Table 4 List of primers

Supplemental Table 5 Putative antiproliferative ISGs

## Acknowledgements

We thank Mayte Suarez-Farinas for advice on statistics and experimental design, and Thomas Moran, Center for Therapeutic Antibody Discovery at the Icahn School of Medicine at Mount Sinai, for kindly providing anti-SARS-CoV Nucleocapsid antibody. We thank Meike Dittmann for A549^ΔSTAT1^ cell lines, Benjamin R. tenOever and Skyler Uhl for technical assistance with plaque assays, and Michael A. Schotsaert for flow cytometry support. We also thank Randy Albrecht for BSL3 facility management and support. We thank Dusan Bogunovic for discussion, helpful advice, and critical reading of the manuscript. We thank all members of the Rosenberg lab for advice and support. This work was supported in part by NIH grants R01 AI151029 and U01 AI150748, and funds from the Department of Microbiology, Icahn School of Medicine at Mount Sinai. Research reported in this paper was also supported by the Office of Research Infrastructure of the National Institutes of Health under award numbers S10OD018522 and S10OD026880. The content is solely the responsibility of the authors and does not necessarily represent the official views of the funders, including the National Institutes of Health.

## Author contribution

Conceptualization: O.D., B.R.R.

Data curation: O.D., B.R.R.

Formal analysis: O.D., R.S.P., B.R.R.

Funding acquisition: B.R.R.

Investigation: O.D., R.S.P., E.J.D., M.R.R.

Methodology: O.D., B.R.R.

Project administration: B.R.R

Supervision: B.R.R.

Validation: O.D., R.S.P.

Visualization: O.D., R.S.P., B.R.R.

Writing – original draft: O.D., R.S.P., B.R.R.

Writing – review & editing: O.D., R.S.P., B.R.R.

## Data availability

Sequence data files from inducible CRISPRa ISG screens are available from the NCBI Sequence Read Archive (SRA), *accession number pending*.

